# A variably imprinted epiallele impacts seed development

**DOI:** 10.1101/339036

**Authors:** Daniela Pignatta, Katherine Novitzky, P.R. V. Satyaki, Mary Gehring

**Affiliations:** Whitehead Institute for Biomedical Research, Cambridge, MA, United States of America; Dept of Biology, Massachusetts Institute of Technology, Cambridge, MA, United States of America

**Author notes:** Corresponding author (MG). These authors contributed equally to this work (Daniela Pignatta, Katherine Novitzky). Current Address: Indigo Agriculture, Charlestown, MA.

## Abstract

The contribution of epigenetic variation to phenotypic variation is unclear. Imprinted genes, because of their strong association with epigenetic modifications, represent an opportunity for the discovery of such phenomena. In mammals and flowering plants, a subset of genes are expressed from only one parental allele in a process called gene imprinting. Imprinting is associated with differential DNA methylation and chromatin modifications between parental alleles. In flowering plants imprinting occurs in a seed tissue – endosperm. Proper endosperm development is essential for the production of viable seeds. We previously showed that in *Arabidopsis thaliana* intraspecific imprinting variation is correlated with naturally occurring DNA methylation polymorphisms. Here, we investigated the mechanisms and function of allele-specific imprinting of the class IV homeodomain-Leucine zipper (HD-ZIP) transcription factor *HDG3*. In imprinted strains, *HDG3* is expressed primarily from the methylated paternally inherited allele. We manipulated the methylation state of endogenous *HDG3* in a non-imprinted strain and demonstrated that methylation of a proximal transposable element is sufficient to promote *HDG3* expression and imprinting. Gain of *HDG3* imprinting was associated with earlier endosperm cellularization and changes in seed weight. These results indicate that epigenetic variation alone is sufficient to explain imprinting variation and demonstrate that epialleles can underlie variation in seed development phenotypes.

**Author Summary:** The contribution of genetic variation to phenotypic variation is well-established. By contrast, it is unknown how frequently epigenetic variation causes differences in organismal phenotypes. Epigenetic information is closely associated with but not encoded in the DNA sequence. In practice, it is challenging to disentangle genetic variation from epigenetic variation, as what appears to be epigenetic variation might have an underlying genetic basis. DNA methylation is one form of epigenetic information. *HDG3* encodes an endosperm specific transcription factor that exists in two states in *A. thaliana* natural populations: methylated and expressed and hypomethylated and repressed. We show that pure epigenetic variation is sufficient to explain expression variation of *HDG3* – a naturally lowly expressed allele can be switched to a higher expressed state by adding DNA methylation. We also show that expression of *HDG3* in strains where it is normally hypomethylated and relatively repressed causes a seed development phenotype. These data indicate that naturally circulating epialleles have consequences for seed phenotypic variation.

## Introduction

DNA methylation is a heritable epigenetic mark that can, on occasion, effect gene transcription and influence development. DNA methylation is a particularly influential regulator of gene expression in endosperm, a triploid extraembryonic seed tissue that supports embryo development. In endosperm, developmentally programmed DNA demethylation causes maternally inherited endosperm genomes to be hypomethylated compared to the paternally inherited endosperm genome (1-3). Methylation differences between maternal and paternal alleles identify their parent-of-origin and establish imprinting, an epigenetic phenomenon in which a gene is expressed primarily from one parental allele (4). Imprinting is theorized to have evolved over conflict between maternally and paternally inherited alleles in offspring over the extent of maternal investment (5,6). Under the kinship theory, silencing of the maternally inherited allele and expression of the paternally inherited allele is predicted to ultimately result for genes where the paternally inherited allele’s optimum expression level in offspring is higher than the maternally inherited allele’s, (7). Comparison of imprinting between species in the Arabidopsis genus has provided empirical support for this hypothesis (8,9).

Recent genomic approaches have revealed extensive natural DNA methylation variation within *Arabidopsis thaliana* (10,11). Whereas the contribution of genetic variation to phenotypic diversity is well-established, the impact of epigenetic variation, or epialleles, on phenotype is only beginning to be understood (12,13). Processes affected by epialleles include patterns of floral development, sex determination, fruit ripening and nutritional content, and senescence, among others (14-19). We previously demonstrated that natural variation in DNA methylation is associated with imprinting variation, with as many as 10% of imprinted genes estimated to be variably imprinted within *A. thaliana* and maize (20,21). Seed development varies extensively among Arabidopsis accessions and has previously been shown to be influenced by parent-of-origin effects (20,22), thus raising the possibility that variation in imprinting could influence seed phenotypes. One of these variably imprinted genes, *HOMEDOMAIN GLABROUS3* (*HDG3*), is a member of the class IV homedomain-Leucine zipper transcription factor (HD-ZIP) family, which regulates diverse aspects of plant patterning and development (23,24). Studies on the function of class IV HD-ZIP genes in trichome differentiation, sepal giant cell formation, and suppression of somatic embryogenesis, among others, have led to the conclusion that class IV HD-ZIP family genes promote endoreduplication and cell differentiation (24-27). Several members of the class IV HD-ZIPs are primarily expressed in endosperm and exhibit imprinted expression patterns, including *FWA/HDG6*, *HDG8, HDG9* and *HDG3* (2,23,28). *FWA, HDG8*, and *HDG9* are maternally expressed imprinted genes (MEGs), whereas *HDG3* is a paternally expressed imprinted gene (PEG) (2,28). The function of the imprinted class IV HD-ZIP genes during seed development, if any, is unknown.

The activity of *HDG3* alleles is correlated with DNA methylation. In endosperm of imprinted strains, the highly expressed paternal *HDG3* allele is methylated and the lowly expressed maternal allele is hypomethylated over a Helitron TE sequence 5’ of the transcriptional start site (2). Maternally inherited endosperm alleles are demethylated by the 5-methylcytosine DNA glycosylase gene *DME*; in *dme* mutants, maternal alleles retain their methylation and are also expressed (2,29). Of 927 Arabidopsis accessions with sufficient methylation data (11), 32 (3.5%) have no methylation in the *HDG3* 5’ region and 871 (94%) have greater than 50% methylation. When strains where *HDG3* methylation is low, such as Cvi or Kz_9, are the paternal parent in crosses with Col, there is no methylation difference between maternal and paternal alleles in endosperm and *HDG3* is biallelically expressed (20). Together, these data suggest that (1) DNA demethylation promotes repression of the maternally-inherited *HDG3* allele whereas DNA methylation promotes expression (or inhibits repression) of the paternal *HDG3* allele and that (2) imprinting variation is due to *cis* epigenetic variation at *HDG3* (20). However, a *cis* or *trans* genetic contribution to imprinting variation cannot be excluded because of DNA sequence polymorphisms between the strains and alleles that do and do not exhibit imprinting.

Here, we show that a naturally occurring epiallele can contribute to variation in seed phenotypes in Arabidopsis. We tested whether *cis* epigenetic variation is sufficient to explain imprinting variation by generating a methylated *HDG3* Cvi allele that mimicked a methylated *HDG3* Col allele. We found that the *HDG3* Cvi allele switched from a hypomethylated, non-imprinted, repressed state to an imprinted, paternally biased, expressed state. Additionally, gain of *HDG3* imprinting altered endosperm development and final seed size. These data indicate that naturally occurring epialleles can have phenotypic consequences in endosperm, a tissue where methylation is dynamic as a programmed part of development.

## Results

### Natural variation in *HDG3* imprinting is associated with gene expression differences

We previously showed that several genes that are imprinted in endosperm when Col is the paternal parent are not imprinted when Cvi is the paternal parent (20). To further examine naturally occurring endosperm gene expression variation, we sequenced the transcriptomes of endosperm from Col x Col and Col x Cvi F_1_ seeds. Comparison of these transcriptomes identified 957 genes that were expressed two-fold or higher in Col x Col and 1187 that were expressed two-fold or higher in Col x Cvi endosperm (Fig 1A; S1 Table). The gene with the lowest expression in Col x Cvi relative to Col is *HDG3*, which is expressed 64-fold lower in Col x Cvi endosperm (Fig 1A).

**Fig 1.**
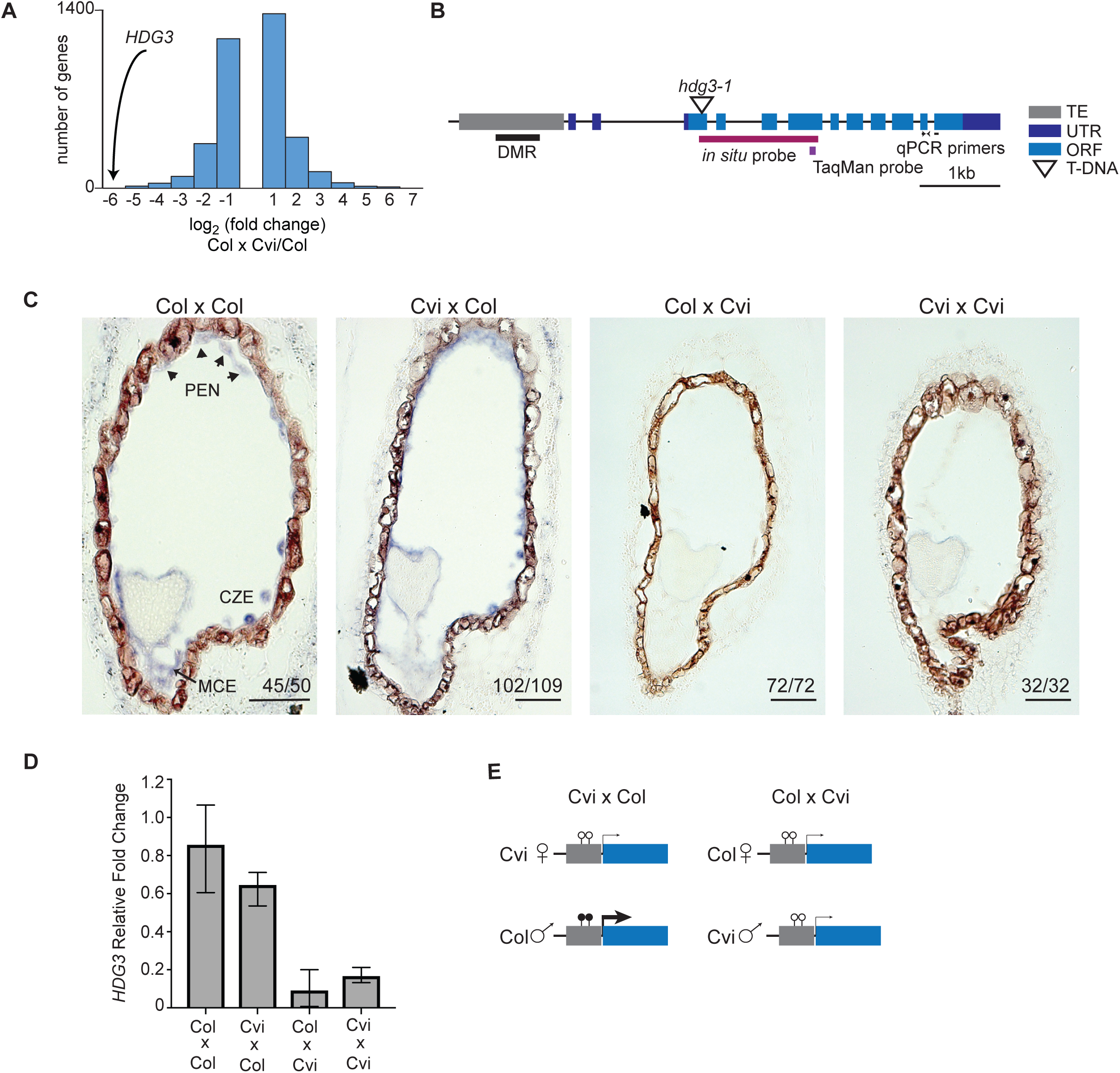
Natural variation in imprinting is associated with differences in *HDG3* expression levels. (A) *HDG3* expression is decreased in Col x Cvi endosperm compared to Col, as determined by mRNA-seq. (B) Schematic of the *HDG3* locus. DMR, differentially methylated region. (C) *In situ* hybridization of *HDG3* (purple) in F_1_ seeds from the indicated crosses. Female parent written first. In crosses where Col is the male parent, *HDG3* is detected in the micropylar (MCE), peripheral (PEN), and chalazal (CZE) endosperm. Arrowheads indicate nuclear-cytoplasmic domains. Number of seeds with shown pattern out of total seeds assayed is in corner of each image. Scale bars, 50 μm. (D) RT-qPCR analysis of relative *HDG3* transcript abundance in F_1_ endosperm. Values are the average of 3 biological replicates, bars represent upper and lower range. (E) Schematic representation of relationship between *HDG3* methylation, expression, and imprinting in endosperm. Thickness of arrows denotes relative expression level. Lollipops represent methylated (filled) and unmethylated (open) cytosines.

We previously reported that *HDG3* is a PEG in Cvi x Col crosses but is biallelically expressed in Col x Cvi (20). To further explore the expression variation of *HDG3*, we performed *in situ* hybridization on developing seeds (Fig 1B-C; S1 Fig). In Col x Col seeds, *HDG3* is expressed specifically in the micropylar, peripheral, and chalazal endosperm, with the highest expression at the heart stage of development (Fig 1C). The same pattern was observed in Cvi x Col (Fig 1C). Whereas *HDG3* expression was detected by in situ hybridization in F_1_ endosperm when Col was the paternal parent, it was not detected in endosperm when Cvi was the paternal parent (Fig 1C). Additionally, we performed RT-qPCR on biological triplicates of Col, Cvi, and Col-Cvi F_1_ endosperm. Expression in Col x Col and Cvi x Col was approximately 10-fold higher than in Cvi x Cvi or Col x Cvi, indicating that *HDG3* expression is higher when it is imprinted (Fig 1D), consistent with the mRNA-seq (Fig 1A) and *in situ* data (Fig 1C). Thus, although expression is from both maternally and paternally inherited alleles in Col x Cvi crosses (and presumably Cvi x Cvi crosses) as detected by mRNA-seq (20), the total expression in those crosses is lower than when *HDG3* is imprinted. As we previously showed that the Cvi allele is naturally hypomethylated (20), together these results suggest that DNA methylation of the *HDG3* 5’ region promotes *HDG3* expression (Fig 1E).

There is also evidence for imprinting variation of *HDG3* in other species. In *Arabidopsis lyrata*, expression of *HDG3* is also specific to the endosperm but levels differ between two accessions, MN47 and Kar, and their reciprocal crosses (S2 Fig). In endosperm with high *HDG3* expression (Kar x MN47), expression is strongly paternally biased (76% paternal instead of the expected 33%), whereas in the reciprocal cross expression of *HDG3* is much lower and more reflective of the 2:1 maternal:paternal ratio in the endosperm (79% maternal) (S2 Fig) (8). The correlation between high expression of *HDG3* and paternal allele bias in *A. lyrata* thus mirrors *A. thaliana*.

### Reduced *HDG3* expression affects seed development

To examine if *HDG3* influenced endosperm development, we compared seeds from *hdg3* mutant plants and segregating wild-type siblings in the Col background. We confirmed predominantly paternal expression of *HDG3* (2,20) by reciprocal crosses between wild type and *hdg3-1* mutants (Fig 2A). When *hdg3* was crossed as a female to a wild-type sibling male, expression of *HDG3* was detected in endosperm in a similar manner as in Col x Col (Fig 2A). In contrast, when wild-type females were crossed to *hdg3-1* mutant males, the accumulation of *HDG3* transcript in endosperm was dramatically affected, with no transcript detected in most cases, despite the presence of a wild-type maternally inherited allele (Fig 2A). We assessed embryo stage and the extent of endosperm cellularization for sectioned wild-type and *hdg3* seeds at 5 days after pollination. Embryo development was more variable in *hdg3*, although this difference was not statistically significant, but endosperm cellularization was significantly delayed compared to wild-type seeds (Fig 2B, S2 and S3 Table). Reciprocal crosses between wild-type and *hdg3* mutant plants indicated that the endosperm cellularization phenotype was dependent on paternal genotype, consistent with *HDG3* function being primarily supplied from the paternally-inherited allele (S2 and S3 Table). Additionally, the weight and area of *hdg3* seeds was slightly reduced compared to Col, suggesting that in the Col background *HDG3* promotes seed growth or filling (Fig 2C-D). Several PEGs have been shown to influence seed abortion phenotypes in interploidy crosses (30,31), but we found no effect of *hdg3* on this process (S3 Fig). To understand the potential molecular consequences of the loss of *hdg3*, we profiled endosperm gene expression in wild-type Col and *hdg3-1* by RNA-seq at 7 days after pollination (DAP) (Fig 3). 150 genes had at least two-fold higher expression upon loss of *hdg3*, while 238 genes had at least two-fold lower expression in *hdg3* mutant endosperm (Fig 3, S4 Table). Differentially expressed genes included developmental regulators such as *Homeobox 3* (*WOX9*) and gibberellin oxidases, which effect the level of a key phytohormone necessary for typical seed development (32) (Fig 3). The loss of *hdg3* also impacted the expression of ten imprinted genes, including the MEG *HDG9* (Fig 3). We hypothesized that the endosperm gene expression phenotypes associated with low expression of *HDG3* from Cvi paternal alleles might in some respects mimic *hdg3* mutants. Indeed, of the 238 genes that are down-regulated in *hdg3* mutants, 100 are also down regulated in Col x Cvi crosses, where *HDG3* expression is also low (Fig 3). This is a highly significant overlap (hypergeometric test in R, p = 6.079e-69) (Fig 3). These data suggest that the Cvi *HDG3* allele, in its hypomethylated and relatively transcriptionally repressed state, could be important for some of the accession-specific developmental traits imparted by Cvi (20,22,33). Thus, to test both the imprinting mechanism and function of *HDG3* further, we introduced methylation at the *HDG3* locus in Cvi.

**Fig 2.**
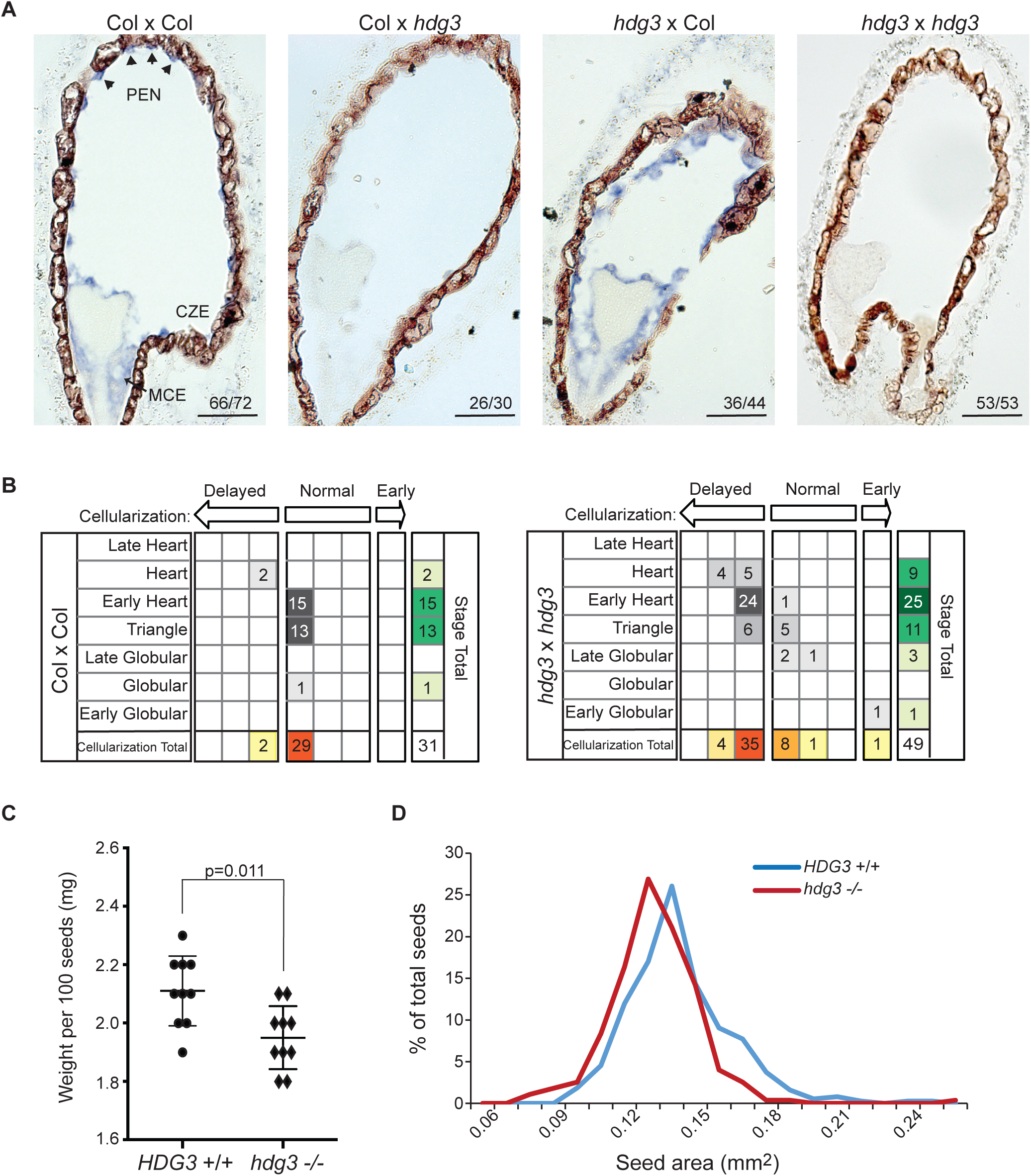
Phenotypic effects of mutation of *HDG3* in Col. (A) *In situ* hybridization of *HDG3* (purple) in seeds from the indicated crosses. Scale bars, 50 μm. (B) Endosperm cellularization is slightly delayed in *hdg3* compared to wild-type at 5 DAP. For each seed the embryo stage was determined and then the stage of endosperm cellularization was defined as normal, early, or delayed given that embryo stage. (C) Seed weight of wild-type and *hdg3* seeds. Individual data points and mean +/-SD shown. P-value from unpaired two-tailed t-test. (D) Seed area is significantly reduced in *hdg3* seed (n=275) compared to wild-type siblings (n=376) (p=8.51e-11 by Welch’s two tailed t-test). Seeds were quantified with ImageJ.

**Fig 3.**
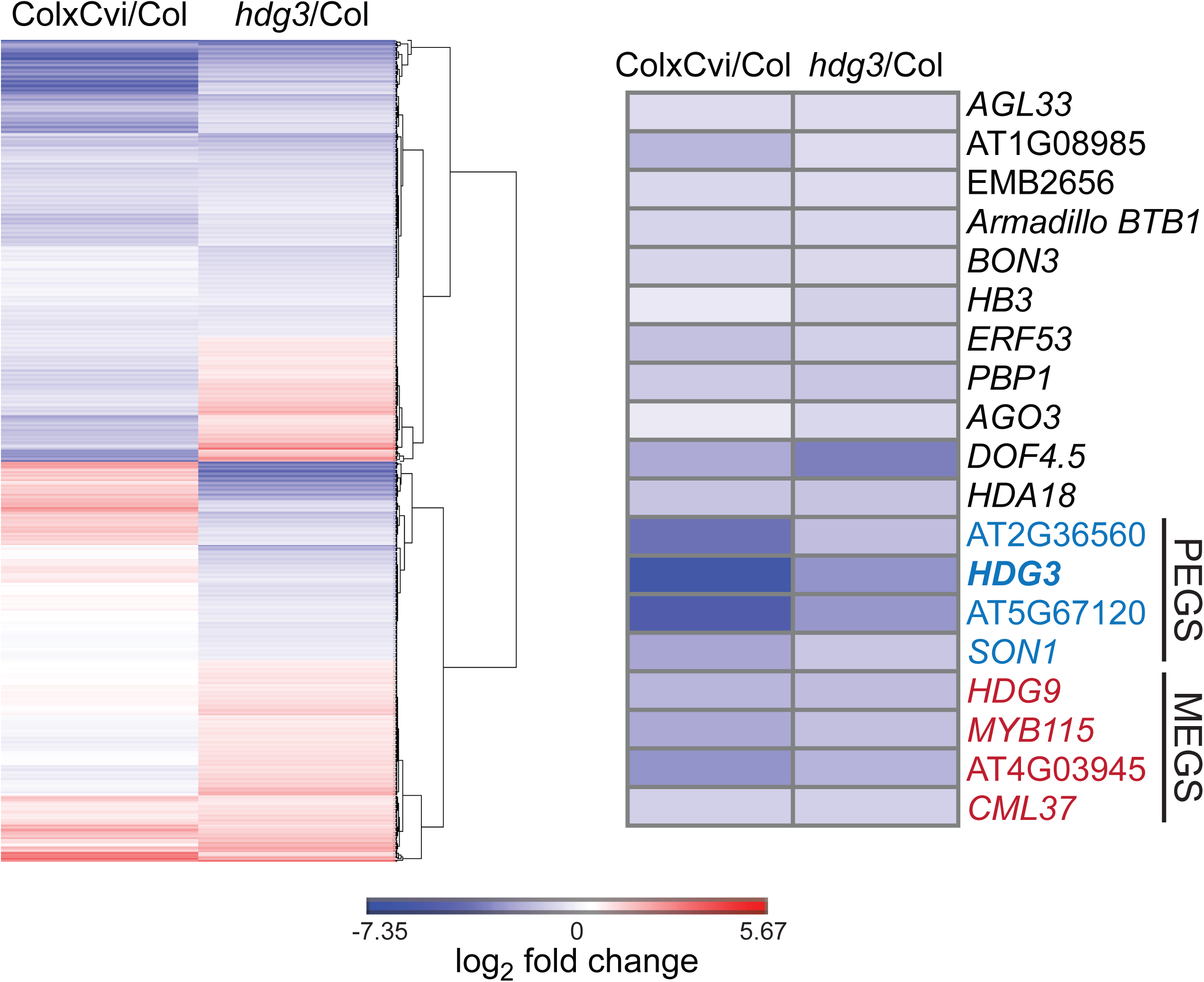
Transcriptional effects associated with low *HDG3* expression. (left panel) Genes downregulated in *hdg3* mutant endosperm also have reduced expression in Col x Cvi endosperm compared to Col. The plot shows the expression profile of genes with significantly altered expression in *hdg3-1* endosperm (p<0.05). Genes were hierarchically clustered by Euclidean distance and complete linkage using Gene-E. (right panel) A subset of putative developmental regulators with reduced expression in *hdg3* and Col x Cvi endosperm.

### An inverted repeat induces methylation in the region 5’ of *HDG3* in Cvi

To distinguish the importance of genetic variation from epigenetic variation for *HDG3* expression and imprinting, we generated transgenic lines in which the endogenous *HDG3* Cvi allele gained methylation in the same region that is methylated in Col. Cvi was transformed with a transgene consisting of an inverted repeat (*HDG3* IR) of the 450 bp *HDG3* 5’ region from Cvi under the control of the constitutive 35S promoter. Processing of the expressed hairpin RNA into small RNAs is expected to direct methylation to the endogenous *HDG3* Cvi locus. We identified multiple independent transgenic lines in which the *HDG3* 5’ region gained methylation in leaves (S4 Fig). DNA methylation was present in the same region as in Col, although non-CG methylation was considerably higher (S4 Fig).

To determine whether the Cvi allele remained methylated when paternally inherited in endosperm, Cvi *HDG3* IR plants from three independent transgenic lines were crossed as males to wild type Col females and DNA methylation was evaluated in F_1_ endosperm by locus-specific bisulfite-PCR. Although the 35S promoter has been reported to have no activity in syncytial endosperm (34), we detected transcripts from the hairpin RNA in endosperm at 7 DAP (S5 Fig). Bisulfite sequencing showed that the paternally inherited *HDG3* Cvi allele from the IR line was hypermethylated relative to the paternally inherited *HDG3* Cvi allele in Col x Cvi endosperm (Fig 4; S6 Fig). The *HDG3* Cvi allele from Col x Cvi *HDG3* IR endosperm was methylated in both CG and non-CG contexts, indicative of RNA-directed DNA methylation, although at a lower level than in leaves or in F_1_ embryos (S6 Fig). Examination of the bisulfite clones indicated some variation in paternal allele methylation, with clones with 0% methylation detected, unlike naturally methylated paternal alleles from Cvi x Col crosses (S6 Fig). This could be due to stochastic silencing of the IR transgene in individual siliques/seeds or ineffective RNA-directed DNA methylation. The maternally inherited Col allele was unaffected in Col x Cvi *HDG3* IR endosperm, remaining hypomethylated like in Col x Cvi endosperm. Thus, we successfully established an alternate epigenetic state specifically for the Cvi *HDG3* allele in endosperm.

**Fig 4.**
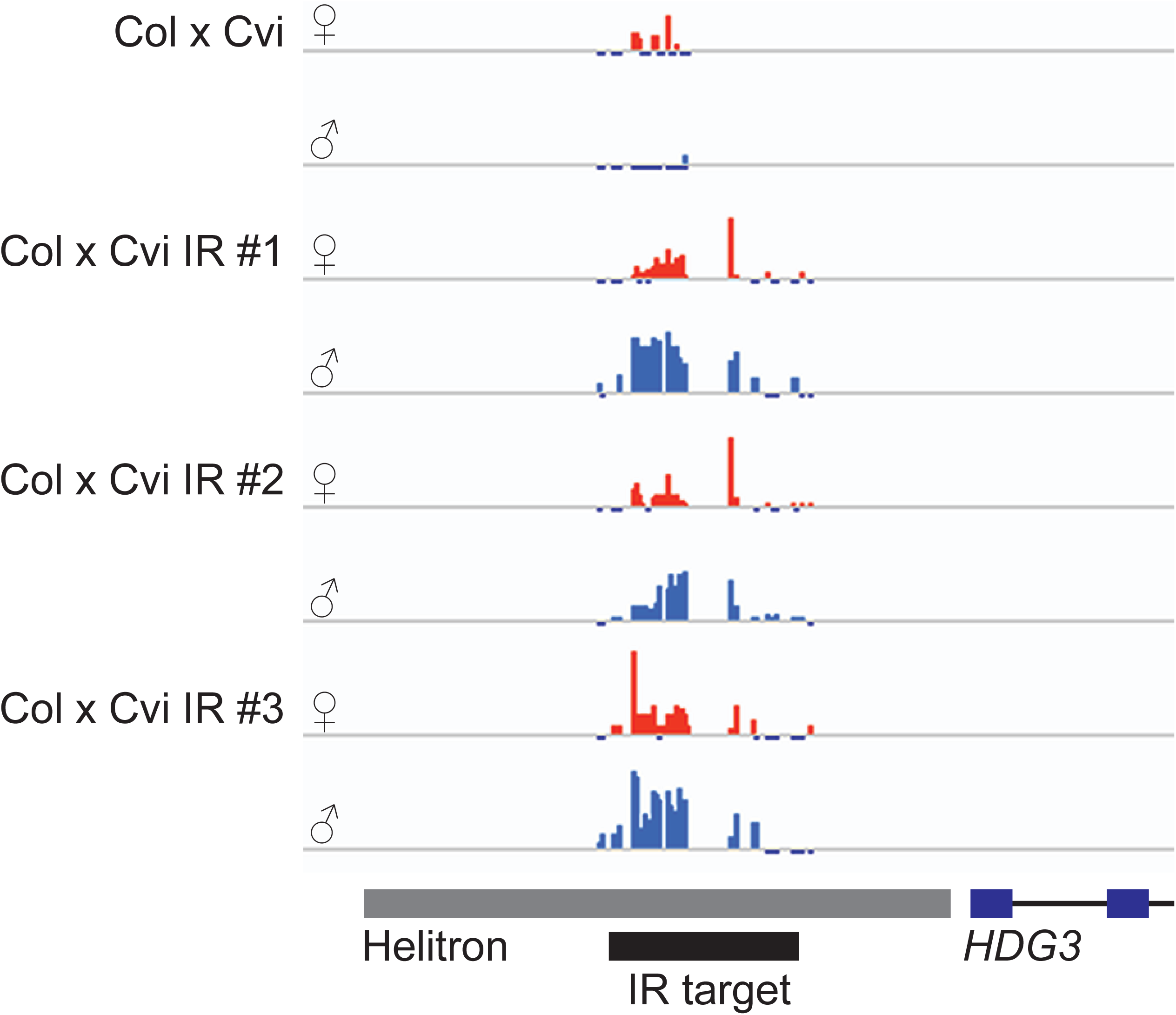
Gain of Cvi *HDG3* paternal allele methylation in endosperm. Total methylcytosine 5’ of *HDG3* in F_1_ endosperm, determined by bisulfite-PCR. Maternally inherited Col allele in orange, paternally inherited Cvi allele in blue. Scale to 100%, tick marks below line indicate unmethylated cytosines. Col x Cvi data published in Pignatta et al., *eLife* 2014.

### Methylation of the *HDG3* 5’ region is sufficient to promote expression and imprinting

Having established a methylated Cvi *HDG3* allele, we tested whether paternal allele methylation was sufficient to switch *HDG3* from a non-imprinted, repressed state to an imprinted, more active state. In two independent lines, *in situ* hybridization of F_1_ seeds from Col x Cvi *HDG3* IR crosses indicated the presence of *HDG3* transcript in endosperm, in contrast to Col x Cvi endosperm (Fig 5A). Hybridization signal was primarily detected in uncellularized endosperm on the chalazal side of the peripheral endosperm (Fig 5A). However, the penetrance of Cvi *HDG3* expression was variable, with about half of the seeds exhibiting *HDG3* expression detectable by *in situ* (Fig 5A). This might be related to the variation in methylation of the *HDG3* Cvi allele in Col x Cvi *HDG3* IR seeds (S6 Fig). Analysis of total *HDG3* transcript abundance by RT-qPCR at 6-7 days after pollination showed that *HDG3* expression was 2-3-fold higher in Col x Cvi *HDG3* IR endosperm compared to Col x Cvi endosperm (Fig 5B). Higher expression of *HDG3* in Col x Cvi *HDG3* IR endosperm is consistent with *HDG3* being more highly expressed when imprinted (Fig 1). Thus, to measure allele-specific expression of *HDG3*, Col and Cvi alleles were distinguished using TaqMan probes in an RT-qPCR assay. In crosses between Col females and three independent Cvi *HDG3* IR lines, the fraction of transcript derived from the Cvi allele increased compared to control crosses between Col females and Cvi males. In Col x Cvi, the Cvi allele accounts for 23% of the transcripts by this assay, in good agreement with prior allele-specific mRNA-seq results (20). In Col x Cvi *HDG3* IR lines, the Cvi fraction was between 50-60%, indicating paternal allele bias (the expectation for non-imprinted genes is 33% paternal) (Fig 5C). This is slightly less than the fraction of paternal allele expression in Cvi x Col crosses by mRNA-seq (79%) (20). Together, these data indicate that the naturally occurring methylation variation at *HDG3* is sufficient to explain imprinting variation. We conclude that the methylated Cvi *HDG3* allele in Cvi *HDG3* IR plants is active and the gene is imprinted.

**Fig 5.**
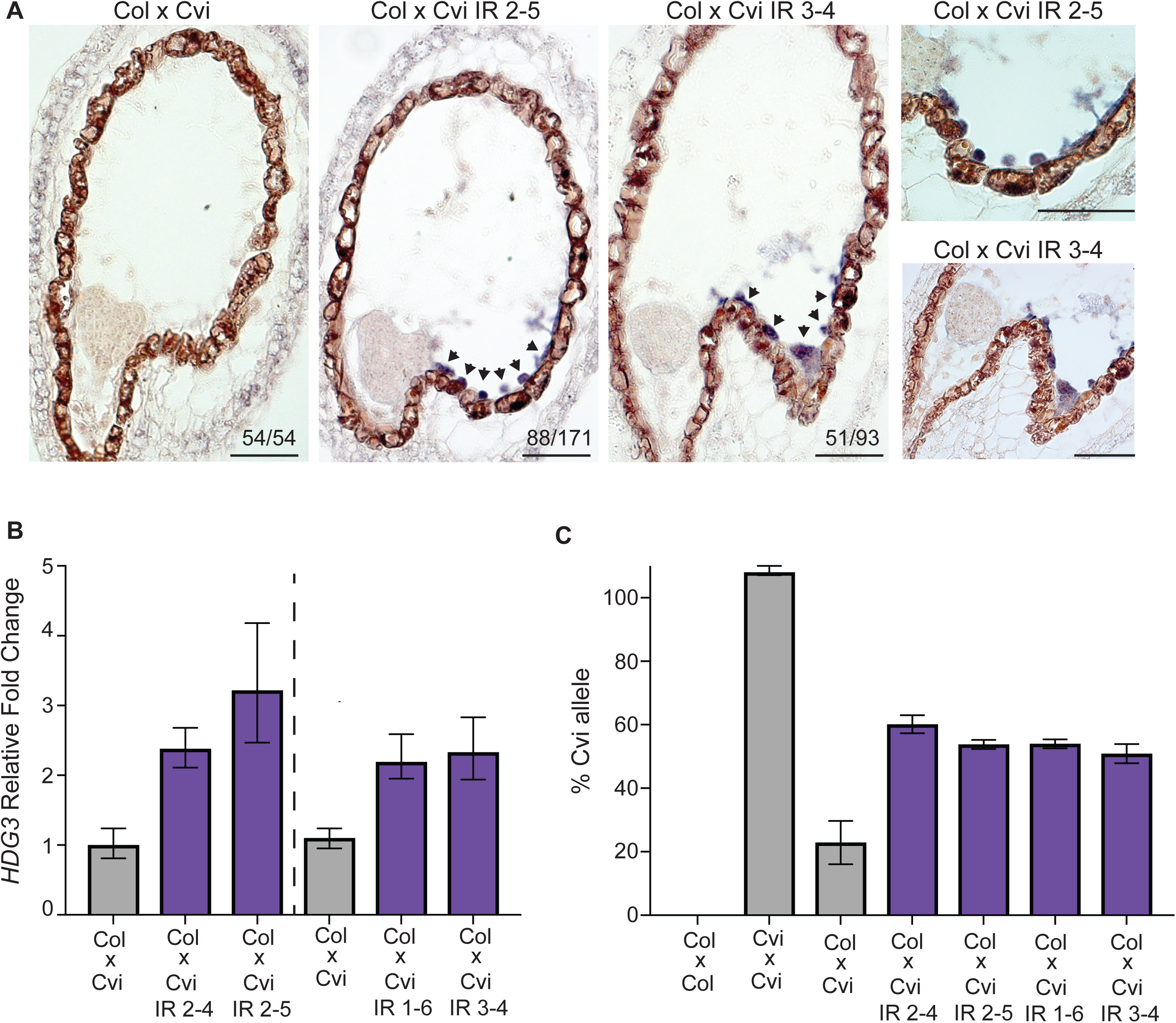
*HDG3* is imprinted in Cvi *HDG3* IR lines. *HDG3 in situ* for indicated genotypes. Arrowheads indicate regions of *in situ* signal. Right panels shows magnification of chalazal region. Scale bars, 50 μm. (B) RT-qPCR of relative *HDG3* transcript abundance in F_1_ endosperm at 6-7 DAP. Dashed line separates experiments done at different times. Left, avg of 3 technical replicates. Right, avg of biological duplicates. Bars show upper and lower range. (C) % of *HDG3* from Cvi allele in endosperm by TaqMan RT-qPCR assay.

### Expression of *HDG3* in Cvi promotes endosperm cellularization

Does the change of *HDG3* expression and imprinting in Cvi affect seed development? To test the phenotypic consequences of expressing *HDG3* from previously repressed Cvi alleles, we compared the phenotypes of seeds from Col x Cvi (low *HDG3* expression) and Col x Cvi *HDG3* IR (2-4-fold increased *HDG3* expression) seeds by sectioning and staining (Fig 6). In crosses with the *HDG3* IR lines, endosperm cellularization occurred at a significantly earlier stage of embryo development, where it was observed as early as the globular stage of embryogenesis (Fig 6A-B, S2 and S3 Table). Whereas endosperm development appeared accelerated, embryo development was significantly delayed (Fig 6B, S2 and S3 Table). The effect on endosperm cellularization was also observed in Cvi x Cvi *HDG3* IR F_1_ seeds, although to a lesser extent (S2, S3 Table). Mature selfed seeds from Cvi *HDG3* IR plants weighed significantly less than selfed seeds from Cvi and had reduced area (Fig 6C-D). This is consistent with known correlations between early endosperm cellularization and the production of smaller seeds (35-37). These observations support the hypothesis that hypomethylation and repression of the Cvi *HDG3* allele is important for Cvi-directed developmental programs and that epiallelic variation contributes to the natural variation in seed development in Arabidopsis.

**Fig 6.**
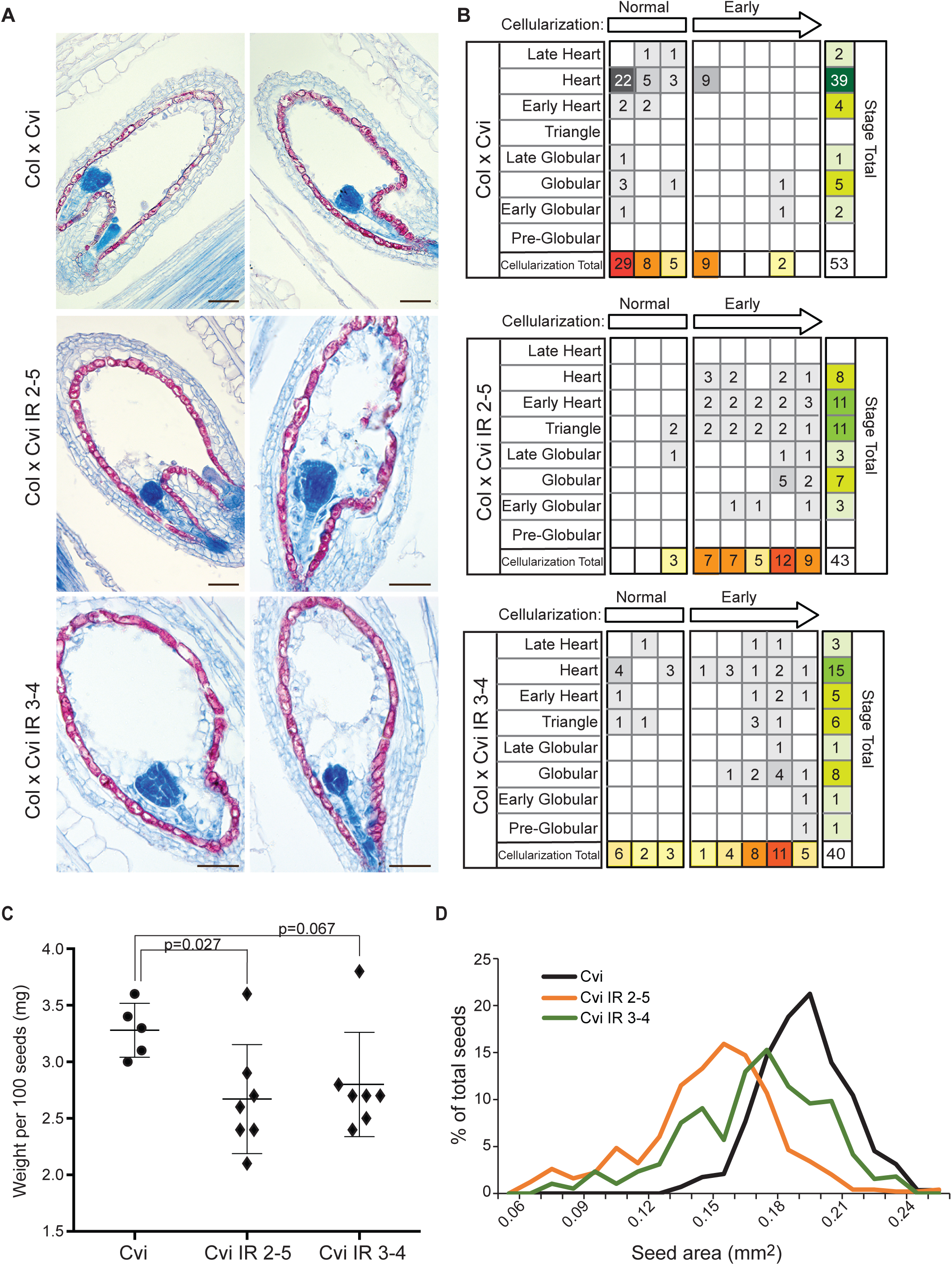
Effects of *HDG3* imprinting on Cvi seed development. (A) Aniline blue and safranin O staining of seed sections at 5 DAP from the indicated F_1_ seeds. Scale bars, 50 μm. (B) Phenotypic characterization of sectioned seeds, assaying degree of endosperm cellularization relative to embryo stage. (C) Seed weight in selfed Cvi and Cvi *HDG3* IR plants. Individual data points and mean +/-SD shown. P-value from unpaired two tailed t-test. (D) Seed area for self-fertilized Cvi (n=287 seeds), Cvi *HDG3* IR 2-5 (n=496) and Cvi *HDG3* IR 3-4 (n=386). Differences between IR seeds and Cvi are significant at p < 2.2e-16 as determined by Welch’s two-tailed t-test.

## Discussion

The establishment, maintenance, and inheritance of DNA methylation are fairly well understood processes. Disruption of methylation patterns by mutations in DNA methyltransferase enzymes have clear gene expression consequences. However, whether or not methylation is regulatory during development – meaning that dynamic loss or gain of methylation is a normal aspect of gene regulation – is less well understood. An exception to this is in the endosperm, where active DNA demethylation in the female gamete before fertilization establishes differential DNA methylation after fertilization, a step that is essential for normal seed development (38). We thus hypothesized that the phenotypic impact of naturally occurring epialleles might be particularly evident in the endosperm, because the differential methylation between maternal and paternal alleles that is required for gene imprinting could be variable across accessions (20). We have shown that *HDG3* represents a case study of this proposed phenomenon. By placing a methylation trigger in Cvi (the *HDG3* IR transgene), we were able to convert the Cvi *HDG3* allele from a hypomethylated to a methylated state. This switch in methylation was sufficient to promote expression of the paternally inherited Cvi *HDG3* allele in endosperm to 3-fold higher levels. Because we altered methylation at the endogenous *HDG3* Cvi locus, which retains all DNA sequence polymorphisms, we have shown that methylation variation alone is sufficient to cause expression, and thus imprinting, variation. However, our results also show that it is unlikely that methylation of the proximal TE accounts for all of the expression differences between paternal Col and Cvi *HDG3* alleles in endosperm. The paternally inherited methylated Cvi allele, while more highly expressed than paternally inherited naturally hypomethylated Cvi allele, was not as highly expressed as paternally inherited methylated Col alleles in endosperm (Figs 1 and 5). Additional *cis* genetic or *trans* genetic or epigenetic variation likely also affects *HDG3* expression levels. Finally, it is not possible to determine from the experiments presented here whether the original difference in methylation between naturally methylated and non-methylated alleles lacks any genetic basis. Cvi lacks the small RNAs associated with the 5’ TE that are found in many other accessions, but the ultimate cause of this difference remains unknown (S4 Fig).

Our experiments also shed light on the relative receptiveness of maternal and paternal endosperm genomes to *de novo* methylation. The *HDG3* inverted repeat transgene should create endosperm small RNAs that are homologous to both Col and Cvi alleles (there are only 4 SNPs and a 3 bp indel between Col and Cvi in the IR target region). Yet, in endosperm from Col x Cvi *HDG3* IR crosses, the paternally inherited Cvi allele had high levels of non-CG methylation, whereas the maternally inherited Col alleles remained hypomethylated despite the presence of the IR transcript (Fig 4, S5 Fig, S6 Fig). In contrast, F_1_ embryos from the same crosses were indeed more highly methylated in the non-CG context on maternal Col alleles compared to maternal Col alleles from Col x Cvi crosses (S6 Fig). Thus, maternally inherited *HDG3* alleles in endosperm are refractory to *de novo* methylation even when a methylation trigger is present, in contrast to maternally inherited *HDG3* alleles in embryos. These results further support findings that once a region is actively demethylated on the maternally inherited endosperm genome, it is “protected” from *de novo* methylation even when triggering small RNAs are present (31).

Finally, although the direct targets of the *HDG3* transcription factor are still unknown, we have shown that natural variation in *HDG3* expression (expressed in Col, low expression in Cvi) has consequences for seed gene expression programs and development (Figs 2, 3, 6 and 7). Expression of *HDG3* in seeds fathered by Cvi caused dramatically early endosperm cellularization and the seeds were smaller and lighter at maturity (Fig 6). These findings are consistent with class IV HD-ZIP genes inhibiting the cell cycle and promoting cellular differentiation (24,27). However, mutation of *hdg3* in Col, while displaying the predicted opposite effect on endosperm cellularization timing, also resulted in smaller seeds weighing slightly less than wild-type (Figs 2, 6 and 7). Although the effects on final seed size are seemingly contradictory and the physiological basis remains incompletely understood, these results are predicted under the aegis of the kinship theory (7). The theory predicts that PEGs promote maternal investment in offspring, which is consistent with the effects of the *hdg3* mutation in Col (i.e. less maternal investment results in smaller seeds). Our results suggest that this effect is specific to a Col seed developmental program. In Cvi endosperm, expression of *HDG3* is seemingly maladaptive, leading to the production of smaller seeds. Cvi naturally produces much larger seeds than Col or L*er*, although fewer in number (20,22,33) (Figs 2 and 6). Our results suggest that the loss of *HDG3* expression in Cvi was an important part of the process that resulted in these phenotypic differences.

**Fig 7.**
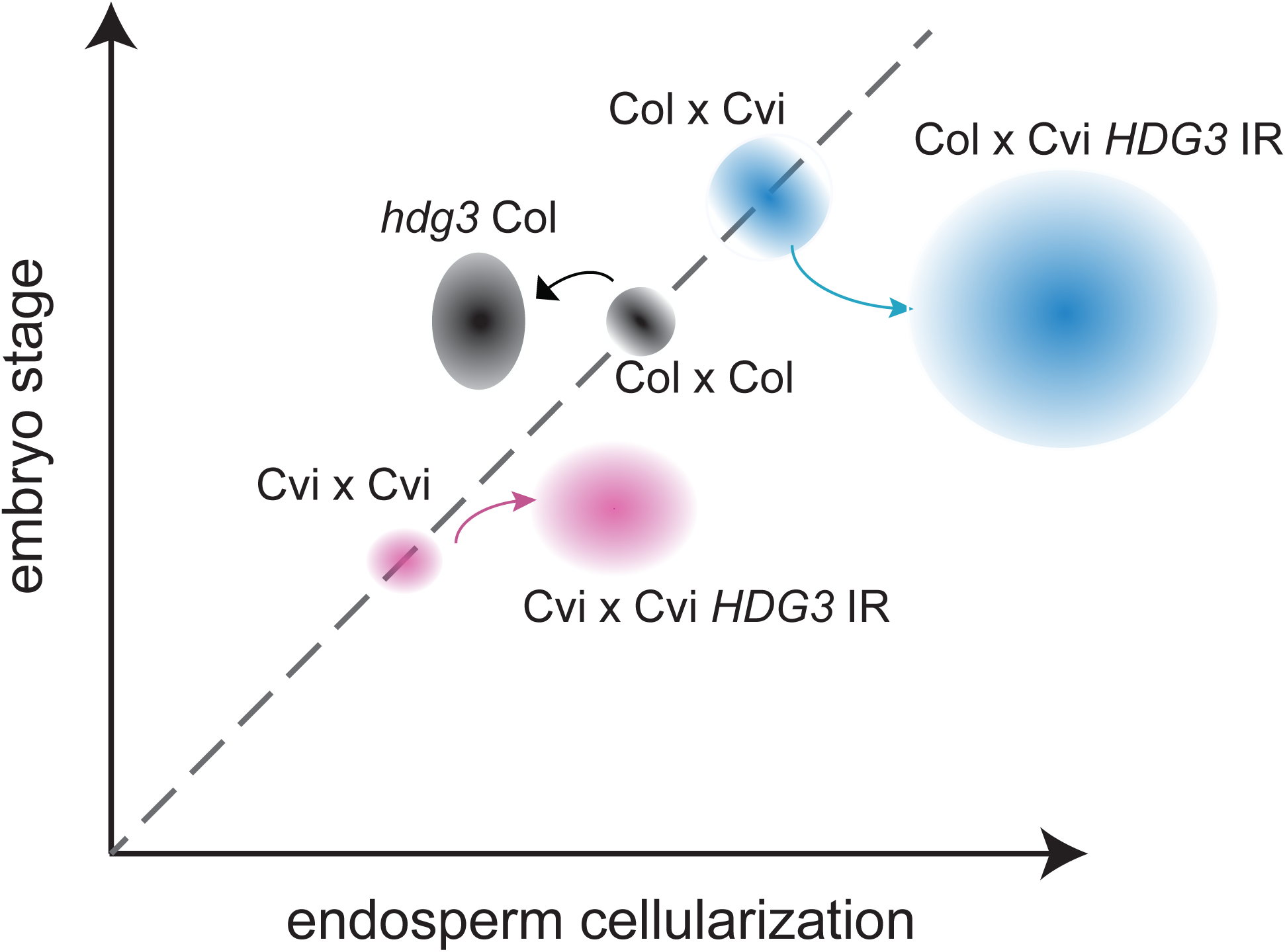
Schematic summary of relative seed development at 5 DAP. Shapes represent phenotypic space occupied by the indicated genotypes.

In summary, we have demonstrated that seed phenotypic differences can be caused by methylation differences at single genes. This study provides further evidence that epigenetic differences underlie developmental adaptations in plants. We have previously shown that the imprinting status of many genes varies between accessions; our current study argues that intraspecific variation in imprinting is an important determinant of seed developmental variation.

## Materials and Methods

### Plant material

The SALK insertion mutant was obtained from the Arabidopsis Biological Resource Center (39). *hdg3-1* (SALK_033462) was backcrossed to Col-0 three times before experimentation. For experiments comparing or crossing wild-type and *hdg3* mutant plants, plants were F_3_ segregants from selfed progeny of *HDG3/hdg3-1*. Plants were grown in a growth chamber or greenhouse with 16-hour days at 22° C. For crosses, flowers were emasculated and then pollinated after 2 days.

### *In situ* hybridization

Controlled floral pollinations were performed for each specified cross. At least two independent *in situ* experiments were performed for each genotype. Siliques were harvested 5 or 6 days after pollination (DAP) and fixed in FAA overnight at 4°C. Following dehydration and clearing (HistoClear, National Diagnostics), samples were embedded in Paraplast Plus (McCormick Scientific), and sectioned at 9 μM (Leica RM 2065 rotary microtome). Ribbons were mounted with DEPC water on ProbeOn Plus slides (Fisher) at 42°C and dried overnight at 37°C. For probes, a 278 bp region of *HDG3* (S5 Table) and previously published 602 bp probe for *PDF1* (40) were amplified from endosperm cDNA and cloned into P-GEM T vectors (Promega). Plasmids containing sense and antisense oriented fragments were identified and linear templates were amplified using M13 forward and reverse primers for probe synthesis. Antisense and sense RNA probes were synthesized in vitro with digoxigenin-UTPs using T7 or SP6 polymerase (DIG RNA labeling kit, Roche/Sigma Aldrich). Probes were subsequently hydrolyzed and dot blots were performed to estimate probe concentration. Pre-hybridization steps were preformed according to (41) except Pronase digestion occurred for 15 minutes at 37°C. Hybridization and post-hybridizations were performed according to (42), with minor modifications. For higher confidence in directly comparing expression patterns, slides corresponding to the cross and its reciprocal were processed face to face in the same pairs for hybridization, antibody, and detection steps. Negative controls consisted of hybridizing sense probes to wild-type tissue and antisense probes to *hdg3* tissue. The sense probe lacked signal (S1 Fig). A probe to *PDF1*, which is expressed in the L1 embryo layer (43), served as a positive control for successful *in situ* hybridization (S1 Fig). Hybridization was performed overnight at 55°C, slides were then washed twice in 0.2X SSC for 60 mins each at 55°C, then twice in NTE for 5 min at 37°C and RNaseA treated for 20 min at 37°C, followed by two more 5 min NTE washes. Slides were incubated at room temperature for 1 hour with Anti-DIG antibody (Roche/Sigma Aldrich) diluted 1:1250 in buffer A (42). Slides were then washed four times for 20 min each at room temperature with buffer A and once for 5 min with detection buffer (42). Colorimetric detections were performed using NBT/BCIP Ready-To-Use Tablets (Roche/Sigma Aldrich) dissolved in water. Slides were allowed to develop 16-24 hours before stopping. Slides were dehydrated, mounted in Permount (Electron Microscopy Sciences) and imaged using a Zeiss Axio Imager M2. Minor level adjustments and smart sharpen were applied to images to compensate for image transfer from live to digital (Adobe Photoshop).

### Seed staining

Plant material was fixed and embedded as previously described and sectioned at 9 μm. Slides were dewaxed twice in xylenes for 5 minutes, rehydrated through a graded ethanol/0.85% salt series from 100%-30%, 1 minute each, stained in 0.6% Safranin O Solution (Cat# 2016-03, Sigma Aldrich) for 5 minutes, washed with water, stained with a saturated 2.5% Aniline blue (Harleco–EMD Millipore, #128-12) in 2% glacial acetic acid aqueous solution for 3 minutes, washed with water, rapidly dehydrated though graded ethanol/salt series to 100%, 5 seconds for each step, and then twice in xylenes for 5 minutes each. Slides were briefly drained, cover slipped and mounted with Cytoseal™ 60 (Thermo Scientific) and imaged using a Zeiss Axio Imager M2.

### Seed phenotypic analysis

Previously processed slides from double staining and *in situ* hybridization experiments were re-examined and used for embryo and endosperm developmental analyses. Using previously published endosperm cellularization and embryogenesis stages (34,44), individual seeds at 5 DAP were scored first for embryo stage and then for respective endosperm stage. Endosperm stage was given a numerical score (−3 to +5) depending on the relative stage of endosperm cellularization compared to the expected endosperm cellularization stage given the embryo stage. Individual seeds with matching embryogenesis and endosperm cellularization stages were scored “normal” and ranged from 0-1; seeds that were scored “early” were defined as being +1.5 to +5 stages further along in the cellularization process compared to normal. Seeds that were scored “delayed” were defined as being −1 to −3 stages behind in the cellularization process compared to normal. To determine whether any developmental differences in endosperm cellularization or embryogenesis were statistically significant, we implemented the asymptotic generalized Pearson chi-squared test from the coin package (45) in R with default scoring weights. Developmental stage was treated as an ordinal variable, while cross genotype was treated as a non-ordinal, nominal variable. Pairwise comparisons were carried out with the R function pairwiseOrdinalIndependence from the rcompanion package. For all tests, embryo development data was collapsed into three categories young (pre-globular to globular), middle (late globular to early heart), and older (heart to torpedo) and detailed endosperm cellularization data was collapsed into the categories delayed, normal, and early.

### Inverted repeat transgene

The 450 bp sequence 5’ of *HDG3* corresponding to a fragment of AT2TE60490 from Chr2: 13740010-13740460 was amplified from Cvi (S5 Table) and cloned into the directional entry vector pENTR-TOPO-D (Invitrogen). The sequence was then inserted twice in an inverted repeat conformation into the vector pFGCGW (46) with a LR clonase reaction (Invitrogen). Cvi plants were transformed with the inverted repeat transgene by floral dipping and T_1_ lines were screened for DNA methylation using bisulfite sequencing. T_3_ plants homozygous for the IR transgene and with a methylated *HDG3* 5’ region in leaves, or their T_4_ progeny, were identified and used for subsequent experiments.

### Quantitative RT-PCR

RNA was isolated from endosperm dissected from seeds at 6 or 7 DAP as described (47) using RNAqueous Micro Kit (Ambion, Life Technologies Corporation). DNAse I-treated RNA (Invitrogen, Life Technologies Corporation) was used for cDNA synthesis with oligo-dT primer using Superscript II reverse transcriptase (Invitrogen) according to manufacturers’ instructions. Quantitative RT-PCR (RT-qPCR) was performed using Fast Sybr-Green mix or TaqMan Mastermix (Applied Biosystems). All reactions were performed in three or four technical replicates using a StepOne Plus Real-Time PCR system (Applied Biosystems). For Sybr-Green based assays, relative expression was calculated using the ddCt method as described (48). The reference gene was AT1G58050 (49). For allele-specific expression in Col-Cvi crosses, a multiplex TaqMan assay was developed by designing primers and PrimeTime^®^ Double-quenched Custom Probes with online tool http://www.idtdna.com/pages/products/gene-expression/custom-qpcr-probes. Cycling conditions were 15 cycles: 95 °C for 15 seconds, 63 °C for 30 seconds, 72 °C for 30 seconds followed by 25 cycles: 95 °C for 15 seconds, 63 °C for 30 seconds with touchdown 0.05°C/cycle, and 72 °C for 30 seconds. The relative expression of each allele within each genotype was calculated using a standard curve (R^2^ value >0.99) as reference. Primer and probe sequences are available in S5Table.

### Bisulfite sequencing

Genomic DNA was isolated from leaves, endosperm, and embryo at 6 or 7 days after pollination using a CTAB procedure. Bisulfite treatment was performed using the MethylCode Bisulfite Conversion Kit (Invitrogen, Life Technologies Corporation) or BisulFlash DNA Bisulfite Conversion Easy Kit (Epigentek Group Inc.) following the manufacturer’s protocols. 2 μl bisulfite treated DNA was used in PCR reactions with 2.5 U ExTaq DNA polymerase (Takara) and 0.4 μM primers using the following cycling conditions (95 °C 3 minutes, 40 cycles of [95 °C for 15 seconds, 50 °C for 30 seconds, 72 °C for 45 seconds], 72 °C for 5 minutes). PCR products were gel purified, cloned using a TOPO-TA (Invitrogen) or CloneJet (Life Technologies) PCR cloning kit and individual colonies were sequenced. Sequences were aligned using SeqMan and methylation was quantified using CyMate (50).

### mRNA-seq

RNA was isolated from endosperm of Col-0, *hdg3-1* and Col-0 x Cvi seeds at 7 DAP as described above. Three replicates for each cross were obtained. DNAse treated RNA was used as input for the SmartSeq Clontech Ultralow RNA-Seq kit. Libraries were constructed by the Genome Technology Core at Whitehead Institute. Six libraries were multiplexed per lane in a Hi-Seq 2500 Standard mode, 40 base, single read run. Each replicate was sequenced to a depth of between 33 and 41 million reads. Reads were processed with Trim_galore using the command “*trim_galore -q 25 --phred64 –fastqc --stringency 5 --length 18”.* Processed reads were aligned to the TAIR10 genome with Tophat2 (51) using the command “*tophat -i 30 -I 3000 --b2-very-sensitive --solexa1.3-quals -p 5 --segment-mismatches 1 --segment-length 18”*. Differential gene expression was detected with Cuffdiff2 (52) and the ARAPORT11 annotation (S1 and S2 Tables). Reads are deposited in GEO GSE118371.

## Acknowledgements

We thank R. Jaenisch and P. Reddien for sharing equipment for *in situ* hybridization, B. Williams for comments on the manuscript, and R. Povilus for assistance with statistical analysis.

## Author Contributions

Conceptualization: MG and DP. Investigation: KN, DP, PRVS, and MG. Writing – Original Draft and Preparation: MG and PRVS. Writing – Reviewing and Editing: KN, DP, PRVS, MG. Funding Acquisition: MG.

## Supporting Information

**Fig. S1.**
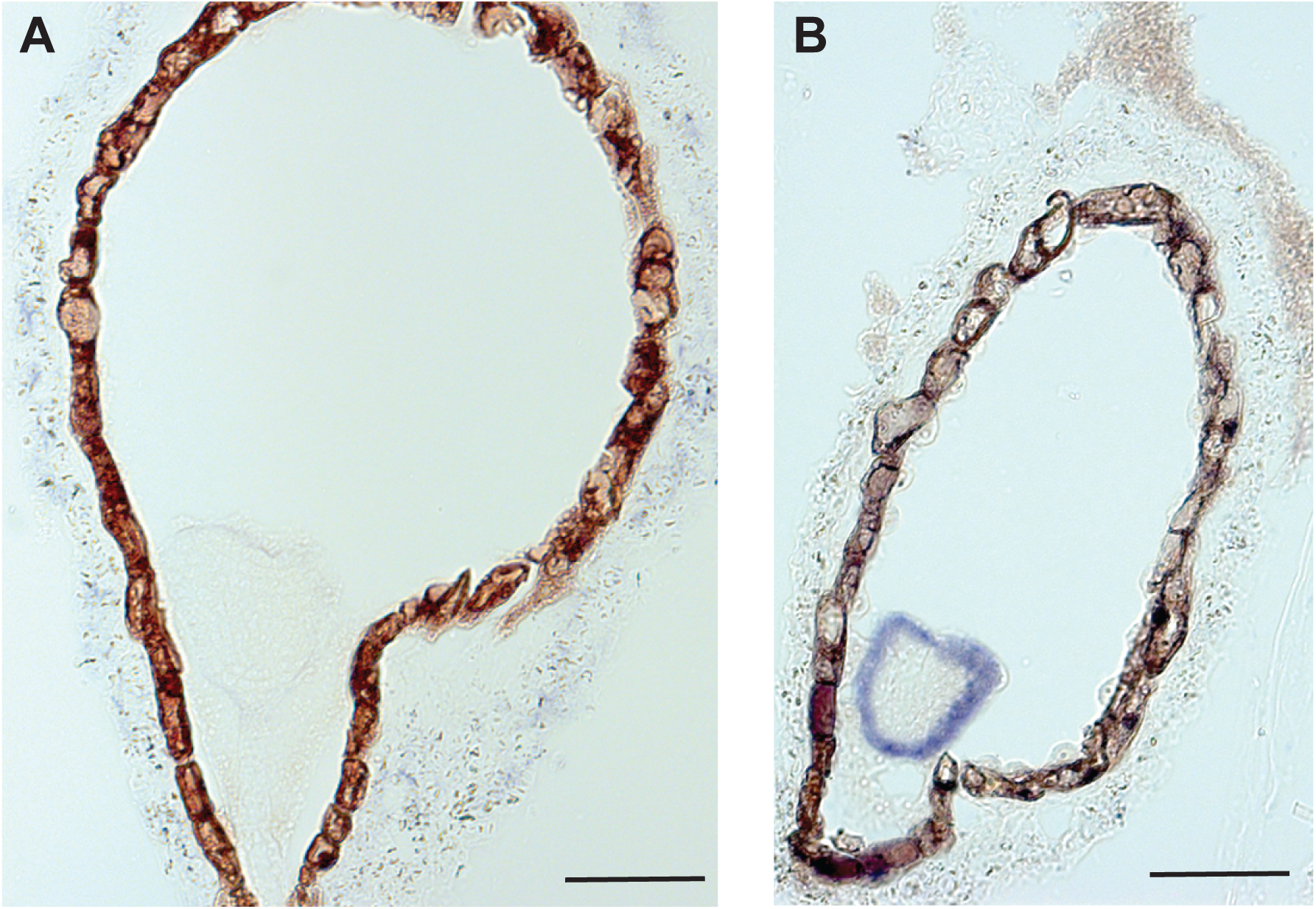
Negative and positive controls for *in situ* hybridization. (A) *HDG3* sense probe hybridization for Col x Col seed. (B) *PDF1* antisense probe hybridization for *hdg3-1* x *hdg3-1* seed. Scale bars, 50 μm.

**Fig. S2.**
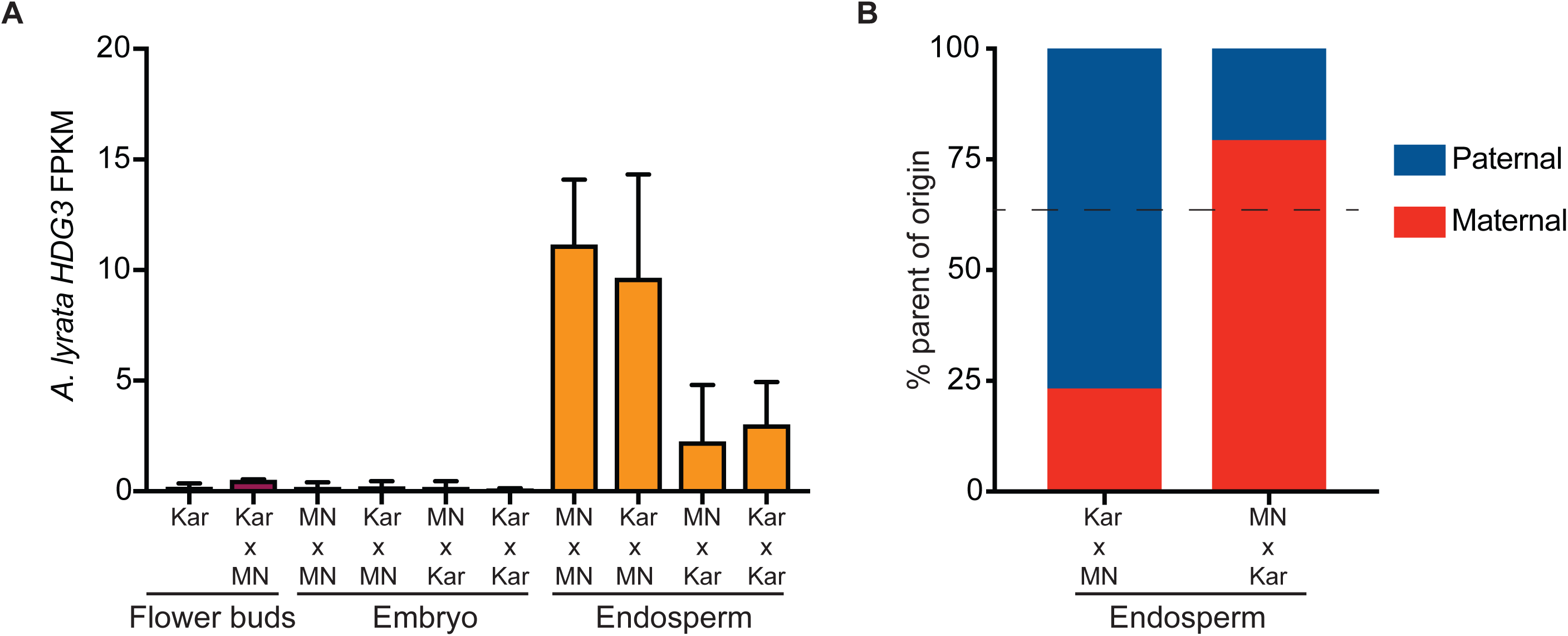
*HDG3* expression in *A. lyrata*. (A) Expression of *HDG3* (ALAl_scaffold_0004_1698 on scaffold 4:15306009-15309421) is specific to endosperm. Bars show mean FPKM values with std deviation for 2-4 biological replicates per genotype. (B) Percent maternal and paternal allele transcripts for the indicated crosses. All data are culled from Klosinska et al., *Nature Plants* 2016.

**Fig. S3.**
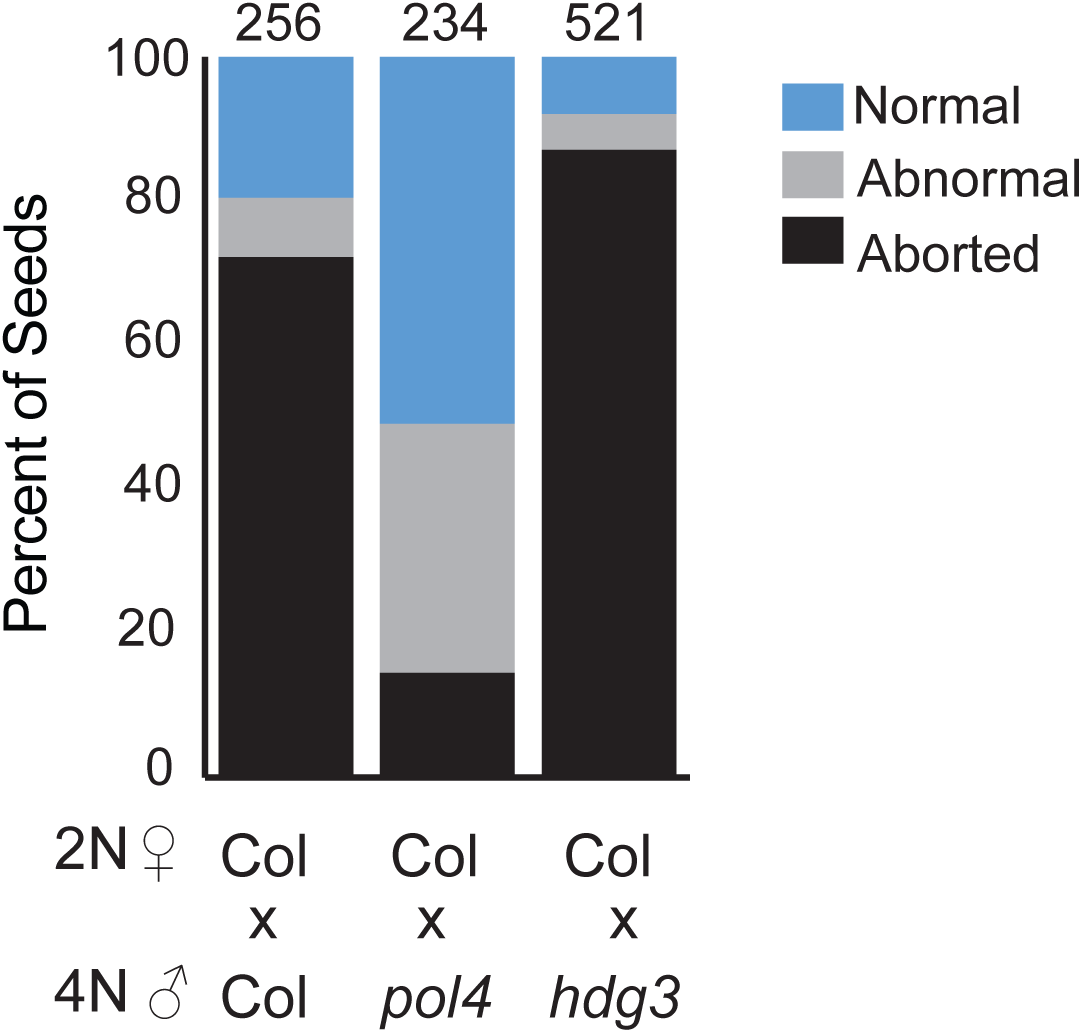
Tetraploid *hdg3-1* does not rescue interploidy seed lethality caused by paternal genomic excess. Tetraploid plants were created by colchine treatment and confirmed by flow cytometry analysis of DNA content. Tetraploid *pol iv* mutants are a positive control for interploidy seed rescue. Number of seeds analyzed on top of bars.

**Fig. S4.**
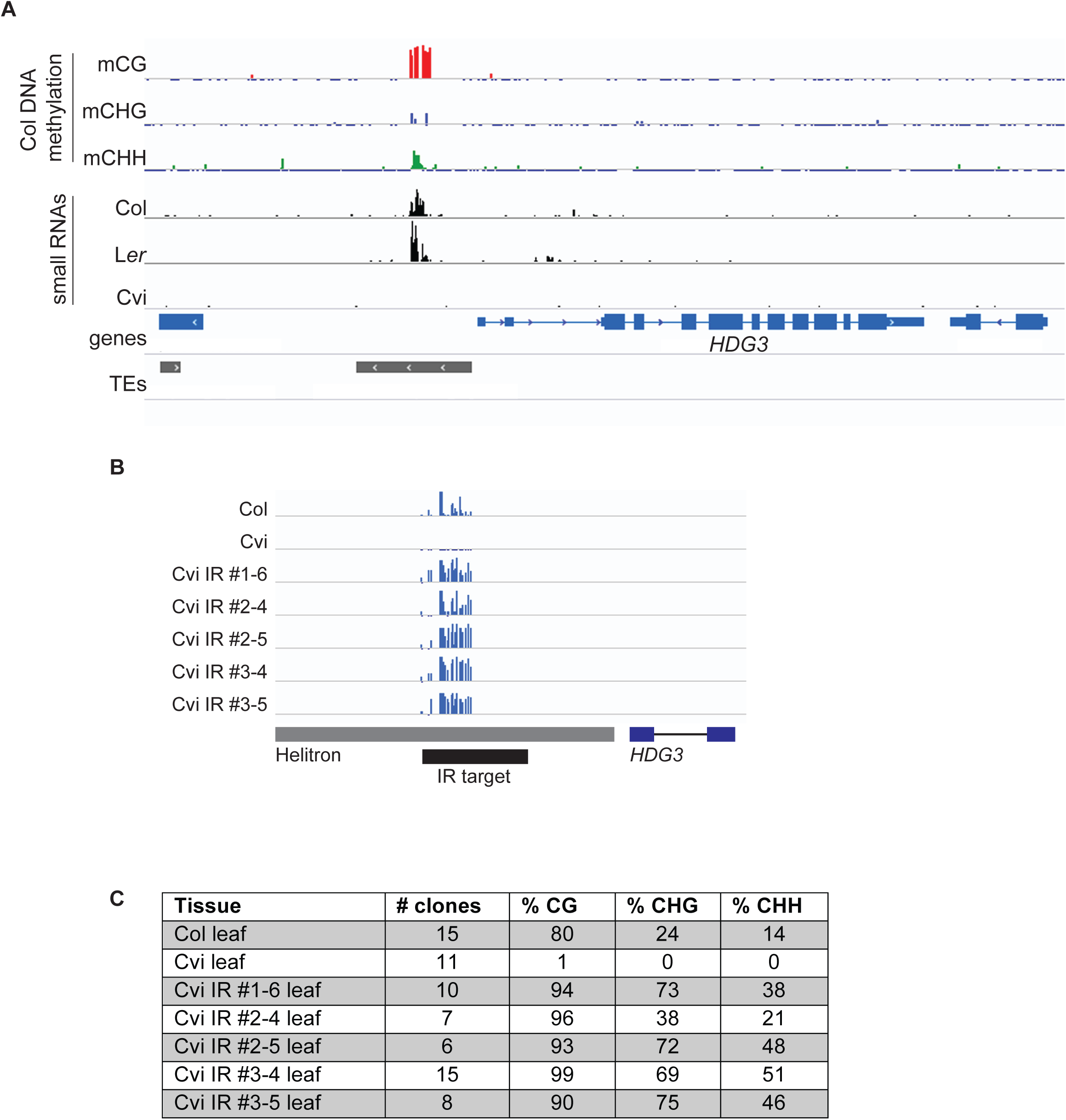
Gain of *HDG3* methylation in Cvi *HDG3* IR lines. (A) Cvi naturally lacks small RNAs at the TE 5’ of *HDG3*. DNA methylation data is from Col embryos (data from Pignatta et al., *eLife* 2014); small RNA data is from Col, L*er* and Cvi embryos (data from Erdmann et al., *Cell Rep.* 2017). (B) Total DNA methylation in Col, Cvi, and Cvi *HDG3* IR leaves as determined by bisulfite-PCR. Scale is 100%, tick marks below line indicate unmethylated cytosines. (C) Quantification of above data. Col and Cvi leaf data are from Pignatta et al., *eLife* 2014.

**Fig. S5.**
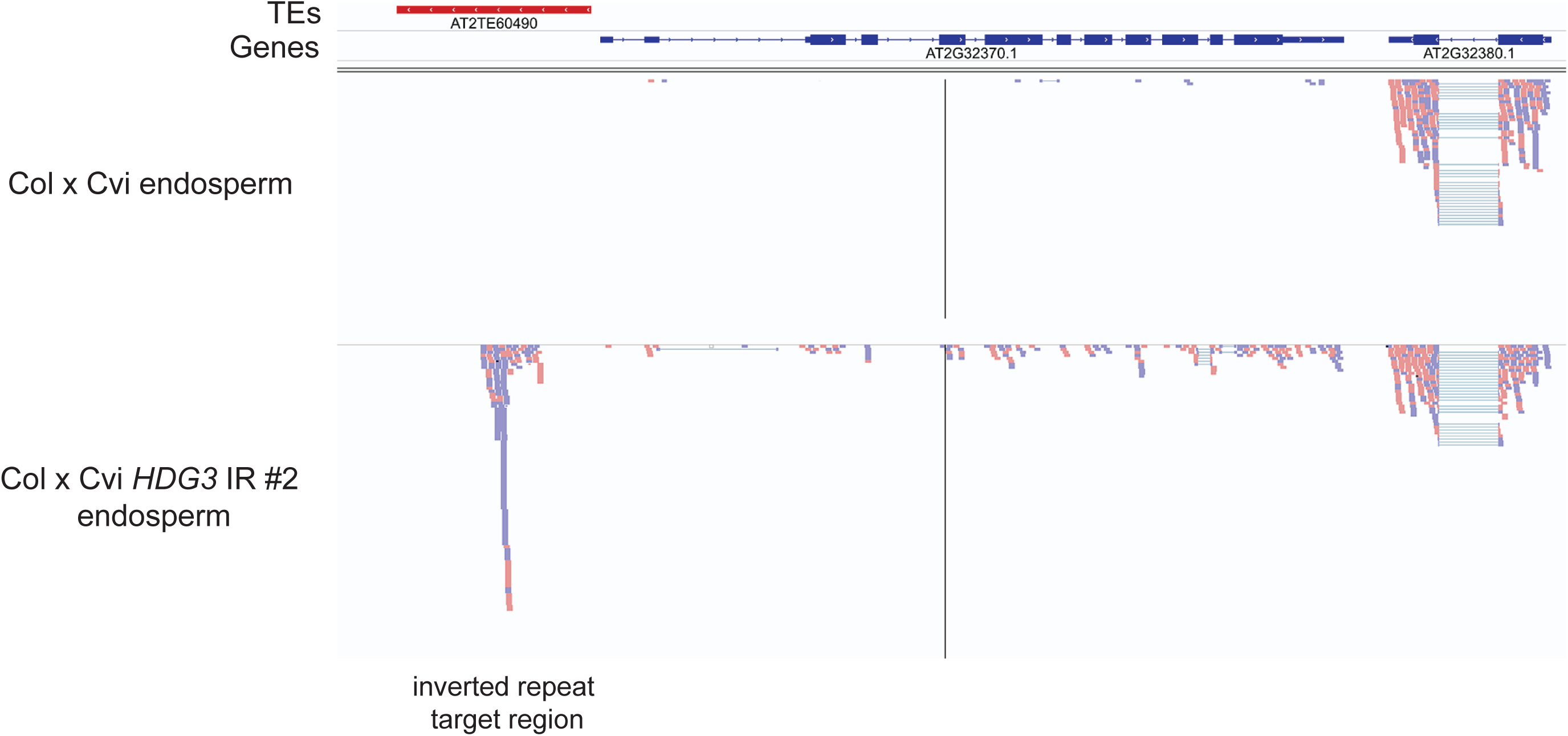
Accumulation of inverted repeat RNA in Col x Cvi *HDG3* IR endosperm. Mapping of mRNA-seq reads to the *HDG3* (AT2G32370) locus in Col x Cvi and Col x Cvi *HDG3* IR #2 endosperm. Reads that match the inverted repeat target region represent expression of the inverted repeat transgene in endosperm from a different location.

**Fig. S6.**
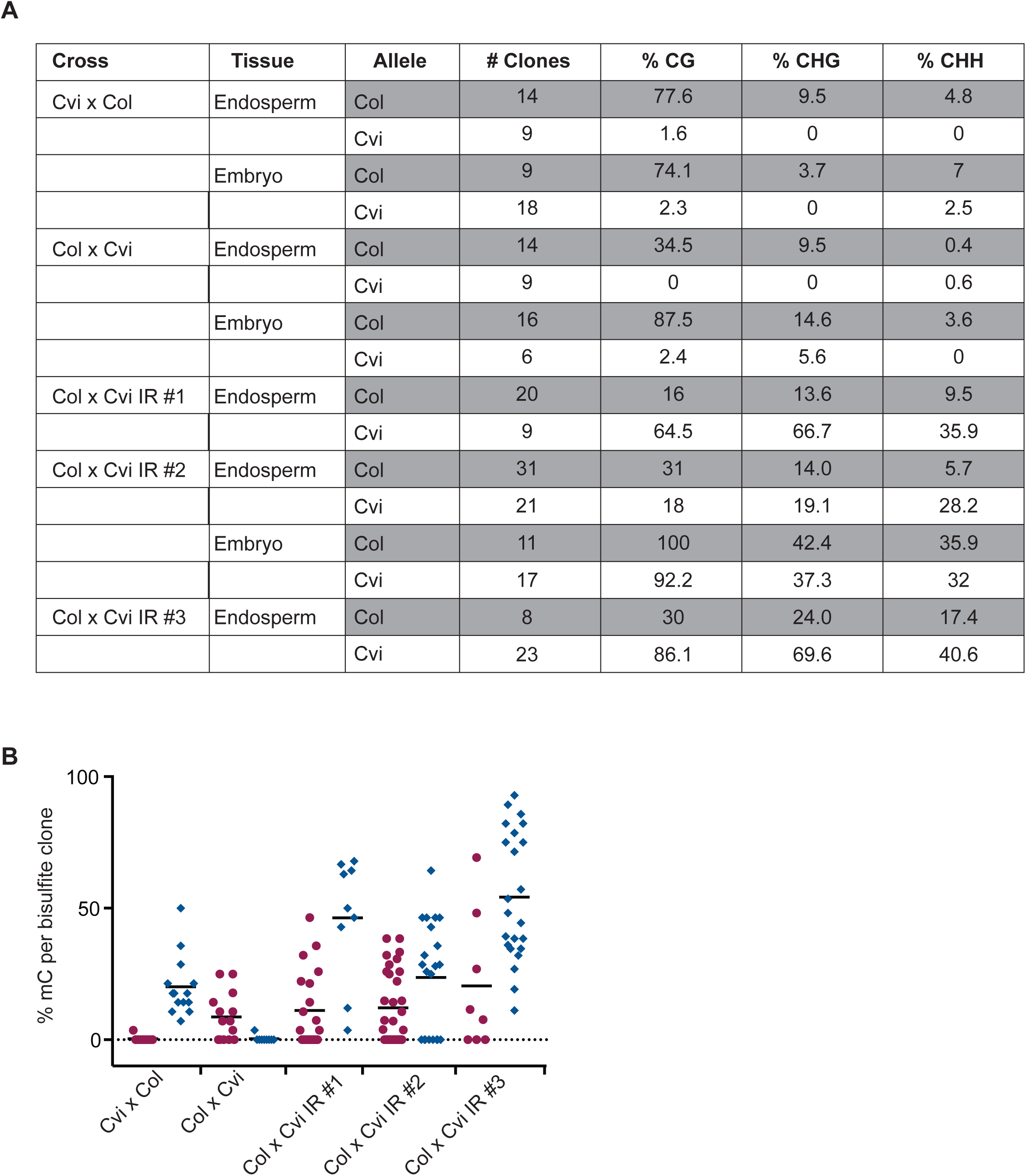
Bisulfite-PCR analysis of *HDG3* methylation in seeds. (A) % methylation from the indicated crosses. Female parent listed first. IR lines are independent transgenic events. Cvi x Col and Col x Cvi data are from Pignatta *et al*., *eLife* 2014. (B) Endosperm total DNA methylation (%) for each bisulfite clone from the above data. Maroon circles, maternal allele clones; blue diamonds, paternal allele clones.

**Table. S1. Gene expression difference in Col x Cvi endosperm vs. Col endosperm at 7 DAP determined using CuffDiff.**

**Table. S2.**
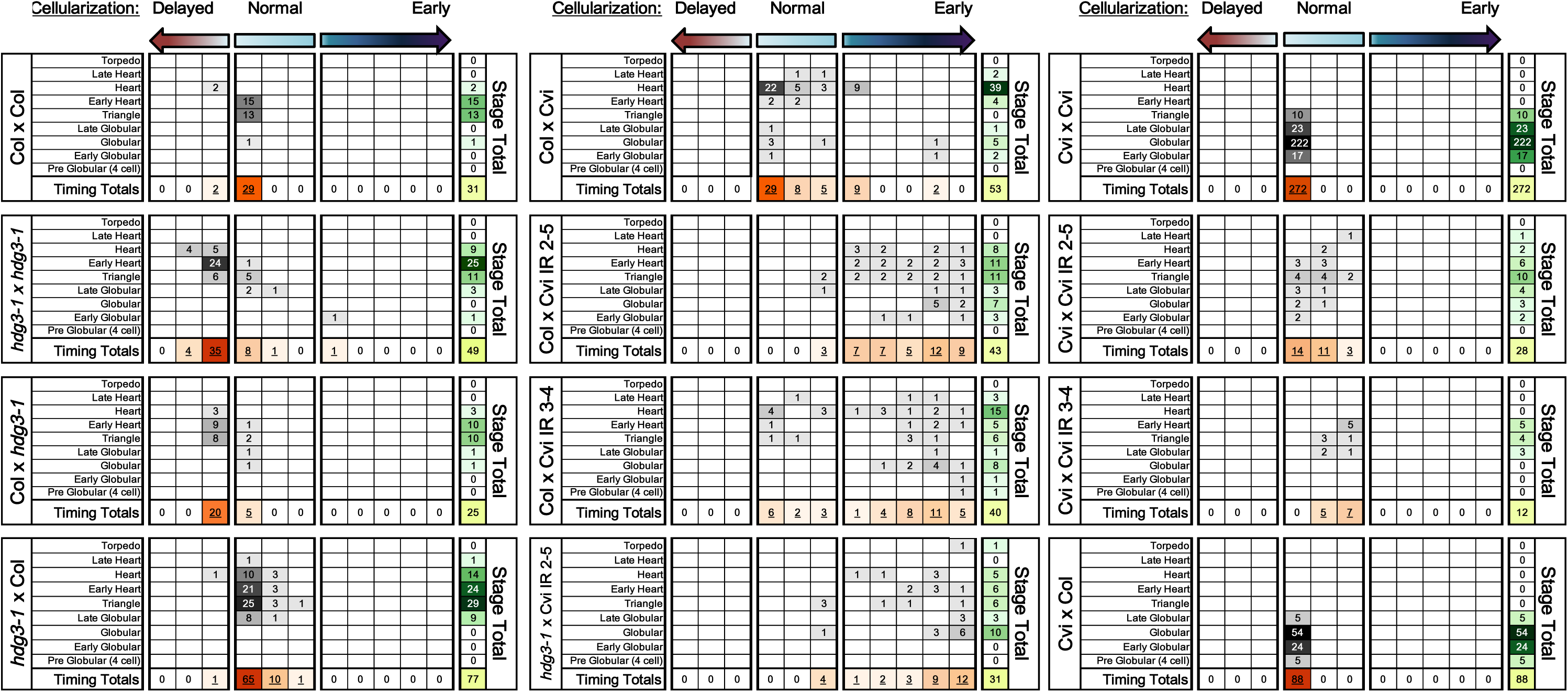
Endosperm cellularization timing and embryogenesis stages at 5 DAP.

**Table. S3.** Tests for statistical significance for pairwise comparisons of differences in endosperm or embryo development.

| Seed Genotype 1 | Seed Genotype 2 | Endosperm diff? | Embryo diff? |
| --- | --- | --- | --- |
| Col x Col | *hdg3* x *hdg3* | *** |  |
| Col x Col | Col x *hdg3* | *** |  |
| Col x Col | *hdg3* x Col |  |  |
| *hdg3* x Col | Col x *hdg3* | *** |  |
| Col x Cvi | Col x Cvi *HDG3* IR 2-5 | *** | *** |
| Col x Cvi | Col x Cvi *HDG3* IR 3-4 | *** | * |
| Col x Cvi | *hdg3* x Cvi *HDG3* IR 2-5 | *** | *** |
| Cvi x Cvi | Cvi x Cvi *HDG3* IR 2-5 |  | *** |
| Cvi x Cvi | Cvi x Cvi *HDG3* IR 3-4 |  | *** |
| Col x Col | Col x Cvi | ** | *** |
| Col x Cvi | Cvi x Col | *** | *** |
| Col x Col | Cvi x Cvi | *** | *** |
| Col x Cvi | Cvi x Cvi | *** | *** |
| Cvi x Col | Cvi x Cvi |  |  |
| Col x Col | Cvi x Col | * | *** |

\*\*\*= difference significant, padj < 0.001; \*\*= difference significant, padj < 0.01; \*=difference significant, padj < 0.05; no stars = no significant difference

**Table. S4. Gene expression differences in *hdg3-1* endosperm vs. Col endosperm at 7 DAP determined using CuffDiff.**

**Table. S5.**
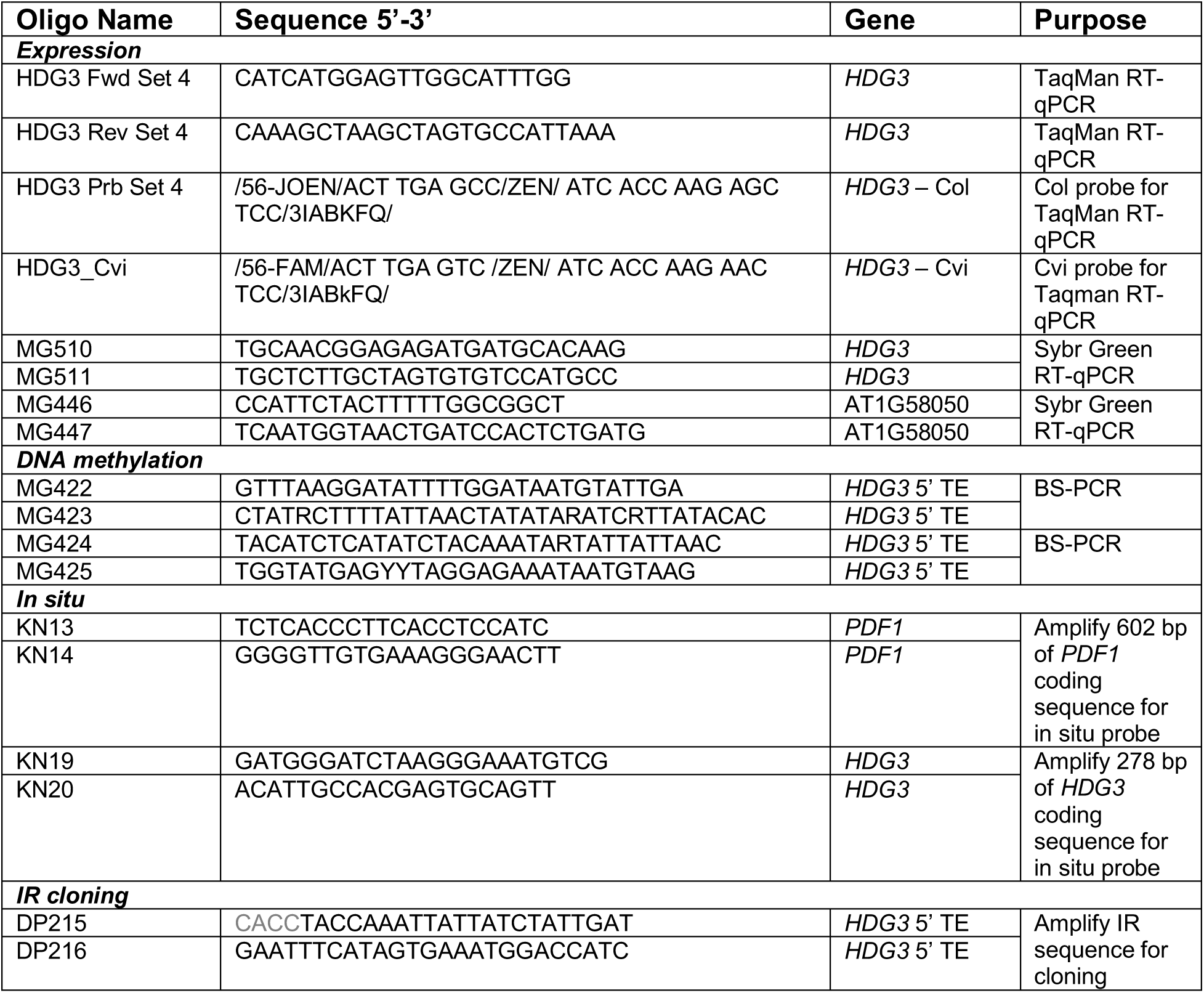
Oligos used in this study.

